# Bayesian Classification of Microbial Communities Based on 16S rRNA Metagenomic Data

**DOI:** 10.1101/340653

**Authors:** Arghavan Bahadorinejad, Ivan Ivanov, Johanna W Lampe, Meredith AJ Hullar, Robert S Chapkin, Ulisses M Braga-Neto

## Abstract

We propose a Bayesian method for the classification of 16S rRNA metagenomic profiles of bacterial abundance, by introducing a Poisson-Dirichlet-Multinomial hierarchical model for the sequencing data, constructing a prior distribution from sample data, calculating the posterior distribution in closed form; and deriving an Optimal Bayesian Classifier (OBC). The proposed algorithm is compared to state-of-the-art classification methods for 16S rRNA metagenomic data, including Random Forests and the phylogeny-based Metaphyl algorithm, for varying sample size, classification difficulty, and dimensionality (number of OTUs), using both synthetic and real metagenomic data sets. The results demonstrate that the proposed OBC method, with either noninformative or constructed priors, is competitive or superior to the other methods. In particular, in the case where the ratio of sample size to dimensionality is small, it was observed that the proposed method can vastly outperform the others.

**Author summary:** Recent studies have highlighted the interplay between host genetics, gut microbes, and colorectal tumor initiation/progression. The characterization of microbial communities using metagenomic profiling has therefore received renewed interest. In this paper, we propose a method for classification, i.e., prediction of different outcomes, based on 16S rRNA metagenomic data. The proposed method employs a Bayesian approach, which is suitable for data sets with small ration of number of available instances to the dimensionality. Results using both synthetic and real metagenomic data show that the proposed method can outperform other state-of-the-art metagenomic classification algorithms.

## Introduction

Many recent studies using both preclinical models and human subjects have highlighted the interplay between host genetics, gut microbes, and colorectal tumor initiation/progression. For instance, the findings in [1] provided novel insights into the complex effects of inflammation on microbial composition/activity and the host’s intestinal mucosa ability to protect itself from microorganisms with genotoxic capabilities. Also, from a mechanistic perspective, the study in [2] demonstrated that Frizzled proteins (receptors that regulate Wnt signaling, required for colonic stem cell maintenance, self-renewal, and repair of the epithelial lining) can be activated by toxin B produced by pathogenic bacteria (*Clostridium difficile*). These findings demonstrate that colonic stem cells are the target of some microbial toxins. Moreover, the results in [3] demonstrated that dietary fiber and fat content have a remarkable effect on the colonic microbiota and its metabolic activity in high-risk vs. low-risk cancer populations.

There is ample evidence that the microbiome (eg, patchy bacterial biofilms) in the gut can modulate host immune cell function, promoting low-grade chronic inflammation. This, combined with intestinal barrier deterioration induced by (1) colorectal cancer–initiating genetic lesions, and (2) crosstalk between microbiota and diets low in fiber, results in the invasion of secreted microbial products (eg, oncotoxins), which can drive tumor growth. Intriguingly, the protective effects of dietary fiber on cancer development may be based on dramatic shifts in microbial community function—particularly short-chain fatty acid production—which stimulate gut mucosal metabolism and increase epithelial barrier function.

The characterization of microbial communities by 16S rRNA gene amplicon sequencing has received renewed interest in the last decade, in part due to the emergence of high-throughput sequencing technology [4]. Microbial metagenomics provides a means to determine what organisms are present without the need for isolation and culturing. Next generation sequencing, applied to microbial metagenomics, has transformed the study of microbial diversity [5]. For amplicons reads it is possible to classify sequence reads against known taxa, and determine a list of those organisms that are present and the read frequency associated with them [6]. In this case an unsupervised strategy can be used to identify proxies to traditional taxonomic units by clustering sequences, so called Operational Taxonomic Units (OTUs).

State-of-the-art classification methods for 16S rRNA metagenomic data include Random Forests (RF) [7, 8] and the phylogeny-based Metaphyl algorithm [9]. RF is a popular classification algorithm, the basic principle of which is to generate a number of random classification trees, and then classify a test point by majority voting among the trees. The most important procedure for generating the ensemble of trees is to search for the best split at each node of the tree over a random selection of from a large number of features, which is the method proposed in [7], but other procedures are possible [8]. Random forests have proved to be very successful in Bioinformatics [19]. In metagenomics, in particular, they have become the industry standard [10, 11]. The Metaphyl algorithm of [9], on the other hand, seeks to introduce prior biological information in the form of microbial species phylogenetic trees, under the assumption that the natural hierarchical grouping of features contained in the phylogeny can help the classification procedure. The Metaphyl classifier is obtained by a regularized maximum-likelihood procedure where the penalty term is guided by the phylogeny tree. Many other approaches for the classification of metagenomic data can be found in the literature; see [9] for a good review of those.

In this paper, we consider a Bayesian paradigm for including prior knowledge, which consists of 1) a hierarchical model for the 16S rRNA metagenomic data, where each microbial sample is represented as a set of operational taxonomic unit (OTU) frequencies; 2) prior distribution construction; 3) posterior distribution calculation from the sample data in closed form; 4) derivation of an *Optimal Bayesian Classifier* (OBC) [12]. The prior distribution on the parameters expresses the biological information available about the problem; in the absence of external knowledge, one may use a “noninformative” prior, or construct the prior from a small portion of the available data (we consider both approaches in our numerical experiments). The hierarchical model proposed here allows computation of the posterior probability distribution in closed form, without a need for computationally-expensive, approximate Monte-Carlo Markov-Chain computations. The posterior distributions are used to derive the OBC, which minimizes the expected error over the space of all classifiers under the assumed likelihood model. Ordinary “Bayes” classifiers minimize the misclassification probability when the underlying distributions are known. However, Optimal Bayesian classification trains a classifier from data assuming the underlying distributions are not known exactly, but are rather part of an uncertainty class of distributions, each having a weight based on the prior and the observed data.

The advantages of the proposed OBC classification algorithm include being applicable: 1) in the absence of phylogenetic information (required by the the Metaphyl algorithm), 2) presence of multiple classes, 3) with small ratios of sample size to dimensionality. The performance of the proposed approach is compared to both the Random Forest and Metaphyl algorithms, as well as a state-of-the-art nonlinear radial-basis function (RBF) Support Vector Machine (SVM) algorithm, using both synthetic and real-world 16S rRNA metagenomic data sets, for varying sample size, dimensionality, and classification difficulty. The results show that the OBC classifier with both noninformative and constructed prior consistently outperform the others on the synthetic data. On the real data sets, the proposed OBC with constructed prior is competitive with the Random Forest algorithm and outperforms the Metaphyl and SVM algorithms on three of the four data sets using pre-defined training-testing splits; the fourth data set is resampled to simulate small ratios of sample size to dimensionality (i.e., number of OTUs), in which case the proposed OBC method, with either noninformative or constructed priors, vastly outperformed all other methods. We remark that a summary of this work, without the real data results or the prior construction method, appeared in [13].

## 1 Materials and Methods

### 1.1 Model for 16S rRNA Metagenomic Data

We propose a Poisson-Dirichlet-Multinomial (PDM) hierarchical Bayesian model for 16S rRNA metagenomic microbial abundance data. Multinomial sampling has been used previously in the study of microbial communities [14, 15]; the novelty of our approach resides in adding Dirichlet and Poisson distributions to allow for OTU sparsity and a varying number of total reads in the sequencing experiment.

Let *M* be the number of OTUs, and let **x** = (*x*_1_,…, *x_M_*) be the microbial abundance vector, where *x_j_* is the number of sequencing reads corresponding to the *j*-th OTU, for *j* = 1,…,*M*. Let 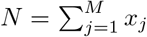 be the total number of sequencing reads. Since *N* is bound to change across different experiments, we model this parameter as a Poisson random variable with parameter λ. The hierarchical model for x assumes a multinomial distribution with probability vector parameter **p** = (*p*_1_,…, *p_M_*) and *N*:

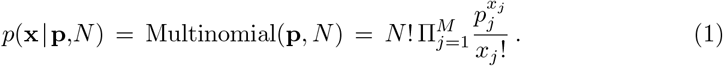

The parameters of the model are **p** and λ. The prior distribution for **p** is assumed to be a Dirichlet distribution with hyperparameter **θ** = (*θ*_1_,…, *θ_M_*), while the prior for λ is a Gamma distribution with hyperparameters *α, β*. We point out that there is in general separate set of parameters and priors, one for each class label (some may be shared between classes), but for simplicity we have not denoted this in this section. See Fig. 1 for a graphical representation of the proposed hierarchical model.

**Fig 1.**
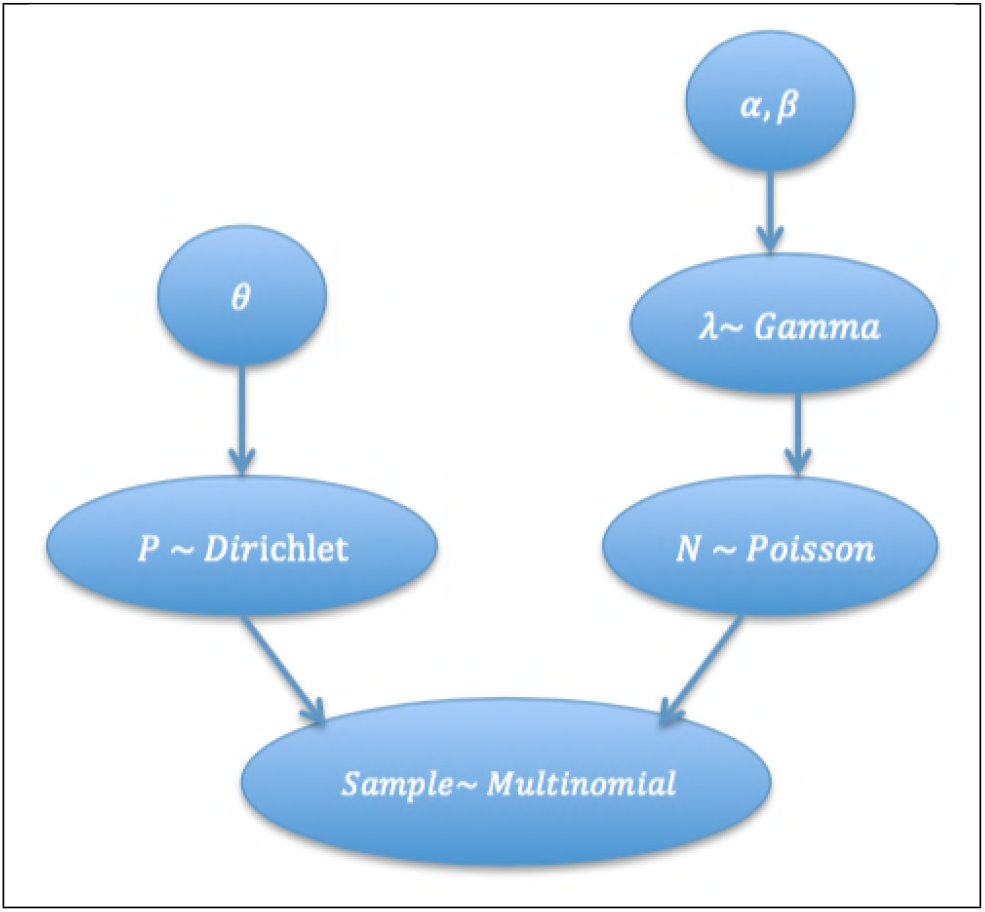
Poisson-Dirichlet-Multinomial hierarchical Bayesian model for 16S rRNA metagenomic microbial abundance data.

### 1.2 Prior Construction

Prior construction concerns setting the values of the hyperparameters to reflect external knowledge about the model. If no external knowledge is available, the hyperparameters may be set to make the priors *noninformative* [16]. In this paper, we assume a noninformative prior for *λ* by setting *α* = *β* =1, but for **p**, we consider both noninformative and informative priors. In the former case, we set ***θ*** = (1,…, 1), in which case the Dirichlet becomes a uniform distribution. In the informative case, we estimate the hyperparameters though a maximum-likelihood procedure using a subset of the available data. This procedure is similar to the one in [17], except that we do not impose any prior knowledge constraints, i.e., the process is entirely data-driven. We describe this procedure next.

We divide the data *S_n_* into two subsamples, one for constructing the prior, and another for deriving the posterior and obtaining the classifier (which is discussed in the next two sections). In all that follows, we consider the binary classification problem, but the approach can be easily adapted to multiple classes (which is the case for some of the data sets considered in Section 2. The class-specific sample sizes 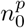 and 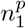 used for prior construction are set to a fixed proportion of the original sample sizes *n_0_* and *n*_1_, respectively, where the indices indicate the class label (in this paper, the ratio is 0.3). In the sequel we omit the class label to simplify the notation, but the Dirichlet parameters are fitted separately for each class.

Let 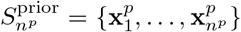 be the sample for constructing the prior, and let 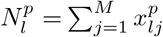 be the total number of sequencing reads in the *l*-th profile, for *l* = 1,…, *n^p^*. Consider the mean log-likelihood

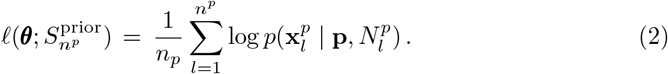

Our objective is to maximize the *expected* mean-likelihood with respect to the parameter to obtain the constructed prior:

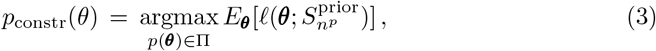

where Π is the set of all Dirichlet priors. Using (1), we have

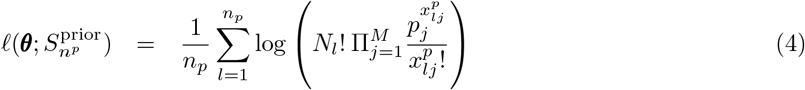

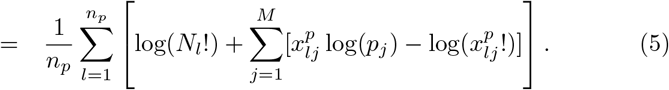

It can be shown that

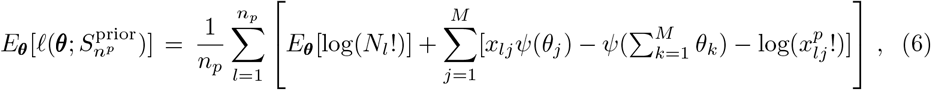

where 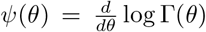 is a digamma function. The optimization in (3) involves transforming the problem to maximization over the hyperparameter space and then applying a numerical procedure, such as conjugate gradient descent, to find the desired hyperparameter values. More details can be found in [17].

### 1.3 Posterior Distribution Calculation

The structure of the model proposed in Section 1.1 allows the exact calculation of the posterior probabilities, avoiding the approximations and computational cost of MCMC methods. Let 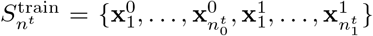 be the training data. The posterior distributions for the parameters (***θ**^y^*, λ) for classes *y* = 0,1 (parameter λ is shared between the classes) are determined as follows:

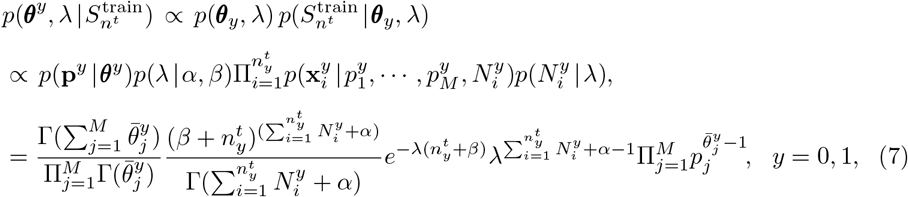

where 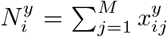 is the number of reads for each profile, 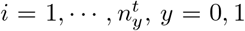, and 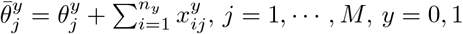.

### 1.4 Optimal Bayesian Classifier

Let *ψ: R^d^* → 0,1 be a classifier designed on training data 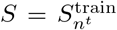, which takes metagenomic abundance profiles ***X*** ∈ *R^d^* into one of the two labels 0 or 1. The error of the classifier is the probability of a mistake given the sample data:

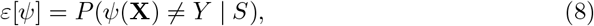

where *Y* ∈ {0,1} denotes the true label corresponding to ***X***.

Now, in the Bayesian setting presently adopted, the joint distribution of **X, *Y***) depends on a random parameter vector ϒ. For example, in our case, ϒ = (***θ***^0^,***θ***^1^,λ).

In addition, the joint distribution depends on the *prevalence c* = *P*(*Y* =1 | ϒ). The expected value of the classification error over the posterior distribution of ϒ becomes the quantity of interest:

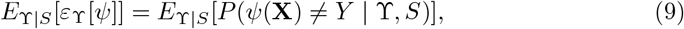

The OBC [12] is the classifier that minimizes the quantity in Eq 9. It was shown in [12] that the OBC is given by

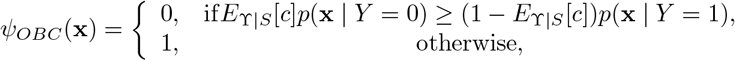

where

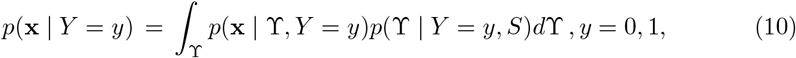

are the *effective class-conditional densities* and *p*(ϒ | *Y* = *y,S*) are the parameter posterior probabilities, for *y* = 0,1.

Plugging Eqs 1 and 7 into 10 leads to an integration that can be accomplished analytically, without a need for MCMC methods, yielding the effective class-conditional densities

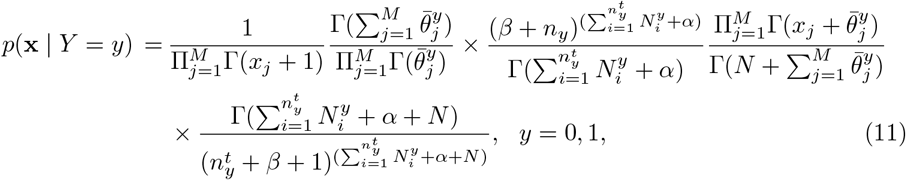

In addition, we assume that the parameter *c* is beta distributed with hyperparameters (*β_0_,β_1_*), independently of ϒ (prior to observing the data). It can be shown that the posterior distribution *p*(*c | S*) is also beta with hyperparameters 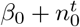 and 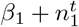, in which case 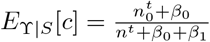. This completes the specification of the OBC classifier in Eq 10.

Fig 2 displays a diagram that summarizes the entire proposed approach to the classification problem.

**Fig 2.**
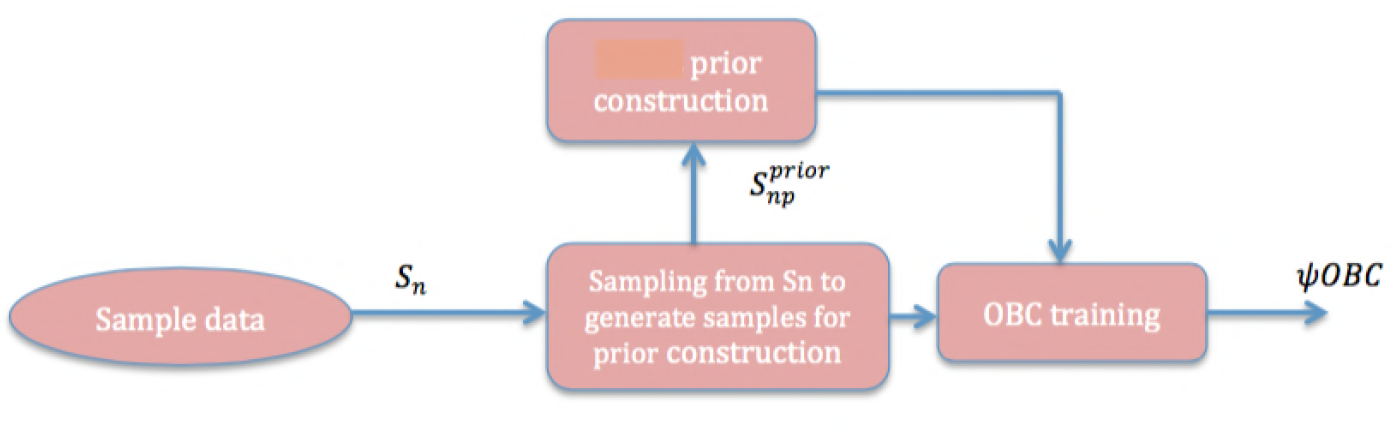
Proposed optimal Bayesian classifier for 16S rRNA metagenomic data.

## 2 Results

In this section, the performance of the proposed classification approach is compared to state-of-the-art metagenomic classification algorithms in the literature, using synthetic and real-world data sets.

## 2.1 Synthetic Data Results

Synthetic OTU abundance data and phylogeny trees were generated using the strategy proposed in [9], which considers a common phylogenetic tree *T* for the OTUs in all the 16S rRNA metagnomic profiles. To generate sample data for each class, the tree is traversed systematically, deciding for each internal node *υ* what fraction of species would come from each of the subtrees rooted at the child nodes of *υ*.

Two parameters are assigned to each node *υ* for each class 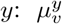 is the average proportion of species that correspond to the subtree rooted at the left child node of *υ* in the class *y*, and *υ* is the variance of this proportion within the class. A new metagenomic profile is generated by sampling the pro-portions of species at each node *υ* according to the normal distribution 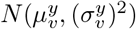. The value of *υ* is in turn sampled from the normal distribution 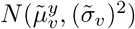, where 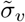 characterizes the variance between the classes, while 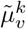 are base values. The within-and between-class variances can be controlled by using the parameters 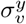 and 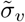, respectively. The exact values of 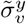 will be sampled at each tree node *σ_υ_* according to 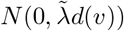 and 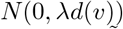, where *d*(*υ*) is the distance between *υ* and the tree root. Note that the parameters lambda and λ influence the difficulty of the classification problem, which is proportional to λ and inversely proportional to 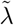.

We consider a total of 128 OTUs and three sample sizes for the training data, *n* = 30, 50, 70, with equal class prevalences, *c* = 0.5. To generate data sets of different complexity, we considered values for 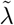 and λ of 0.5, 1 and 1.5. The sample sizes *n*_0_ and *n*_1_ are fixed at *n*_0_ = *n*_1_ = *n*/2. Classifier accuracy is obtained by testing each designed classifier on a large synthetic test data set, and averaging the results over 1000 iterations using different synthetic training data sets.

Fig. 3 compares the performance of our proposed optimal Bayesian classifier, using both noninformative and constructed priors, against that of the kernel SVM [18], RF [19] and MetaPhyl [9] classification algorithms, mentioned in Section 1. As expected, the classification error rate is reduced when the variance within class was decreased and variance between classes was increased. Also as the sample size increases, classification performance improves for all classification rules. We observe that the kernel SVM classifier clearly performs poorly comparing to other classifiers, the RF and Metaphyl classifier have comparable performance overall, while the proposed OBC classifier exhibits the best performance over different values for between and within class variances.

**Fig 3.**
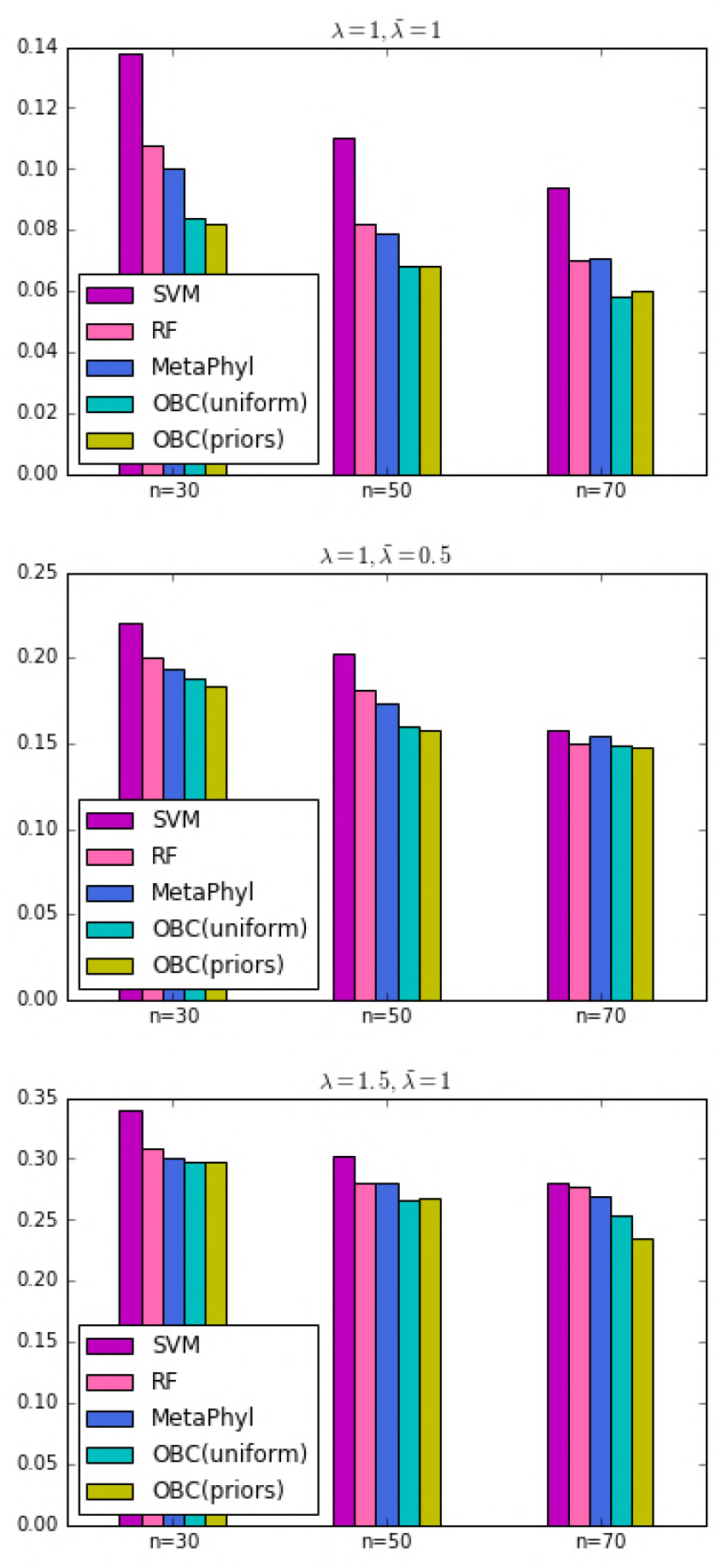
Comparison of Kernel SVM, RF, MetaPhyl and Optimal Bayesian classifier with uniform and constructed priors on simulated data sets for varying sample size and within- and between-class variances, reflecting an increasing difficulty of classification. Each sample abundance profile contains 128 OTUs.

## 2.2 Real Data Results

Here, we consider four different 16S rRNA metagenomic data sets. The sample sizes, dimensionality, and number of classes of each data set are displayed in Table 1. Each of these data sets is accompanied by a phylogeny tree, which allows the application of the Metaphyl classification algorithm.

**Table 1.** Summary of real metagenomic data sets

| Data set | Training samples | Test samples | Number of OTUs | Number of classes |
| --- | --- | --- | --- | --- |
| CBH | 415 | 207 | 2741 | 6 |
| FS | 68 | 33 | 565 | 3 |
| FSH | 68 | 33 | 565 | 6 |
| Twin Gut | 257 | — | 5462 | 2 |

### Costello et al. Body Habitats (CBH) [20]

The biogeography of bacterial communities on the human body is critical for establishing healthy baselines and it helps to detect differences associated with diseases. These data included sample communities from six major categories of habitat: External Auditory Canal (EAC), Gut, Hair, Nostril, Oral cavity, and Skin. This data set is an example of a relatively easy classification task due to the generally pronounced differences between the communities. These microbiota, varies systematically across body habitats and time; such trends may reveal how microbiome changes cause or prevent disease [20].

### Fierer et al. Subject (FS) [21]

In this work they show that skin-associated bacteria can be readily recovered from surfaces and the structure of communities can be used to differentiate objects handled by different individuals. This data set contains all samples from the ‘‘keyboard” data set, for which at least 397 raw sequences were recovered [14]. The class labels are the anonymized identities of the three experimental subjects. This classification task is the easiest of all four data sets because of the clear distinctions between the individuals, the fact that all of the samples come from approximately the same time point, and the large number of training samples available for each class.

### Fierer et al. Subject Hand (FSH) [22]

The influence of sex and washing on the diversity of hand surface bacteria is studied in this work. Data set contains the palmar surfaces of the dominant and non-dominant hands of 51 healthy young adult volunteers to characterize bacterial diversity on hands and to assess its variability within and between individuals. This data set is a more challenging version of the previous ones. The class labels are the concatenation of the experimental subject identities and the label of which hand (left vs. right) the sample came from on that individual. There were three subjects, and so there are six classes in this data set.

### Turnbaugh et al. Twin Gut [23]

This data set contains the faecal microbial communities of adult female monozygotic and dizygotic twin pairs concordant for leanness or obesity, and their mothers, to address how host genotype, environmental exposure and host adiposity influence the gut microbiome. This data set is a challenging classification task because the classes correspond to microbial communities from the same body habitat and thus are very similar.

We compare our classifier with RF, kernel SVM and MetaPhyl classifiers, described in Section 1. For comparing different methods, we used the training and test data splits provided in the CBH, FSH, FS studies. For the Twin Gut data set, we iteratively sampled subsets of data of small sample sizes from the data set for training, applied the classification rules, and tested the resulting classifiers on the remaining large collection of points not selected. This was done to simulate the effects of sample sizes on classifier training. This process is repeated 500 times and the results are averaged. In all cases, for the OBC with prior construction, the “training” data includes both the subsample for prior construction and for the subsample for training proper (i.e., posterior inference).

The classification error rates for the CBH, FSH, and FS data sets are displayed in Table 2. We can observe that the proposed OBC with constructed priors outperforms all others on the CBH and FSH data sets, while being slightly outperformed by the state-of-the-art Random Forest classifier on the FS data set; in fact, the error rates for all classifiers is small on this latter data set, revealing an easy classification problem. Notice that the results for the OBC with constructed priors are always better than the OBC with noninformative priors, as expected.

**Table 2.** Classification error rates for the CBH, FS, and FSH data sets

| Method | CBH | FS | FSH |
| --- | --- | --- | --- |
| RF | 0.099 | **0** | 0.026 |
| SVM | 0.133 | 0.081 | 0.041 |
| OBC | 0.116 | 0.034 | 0.031 |
| OBC(constructed priors) | **0.075** | 0.015 | **0.021** |
| MetaPhyl | 0.104 | 0.022 | 0.036 |

The results for the Twin Gut data sets, for varying sizes of the training data set are displayed in Figure 4. Here the small-sample effects become clear. In particular, it is evident that Random Forests cannot handle very small training sample sizes well. The performance of the proposed method is vastly superior to the others fon this data set. We believe that the reason is that the ration between sample size and dimensionality (i.e., the number of OTUs) is much smaller in this data set than the others. In addition, the classes are unbalanced in this data set.

**Fig 4.**
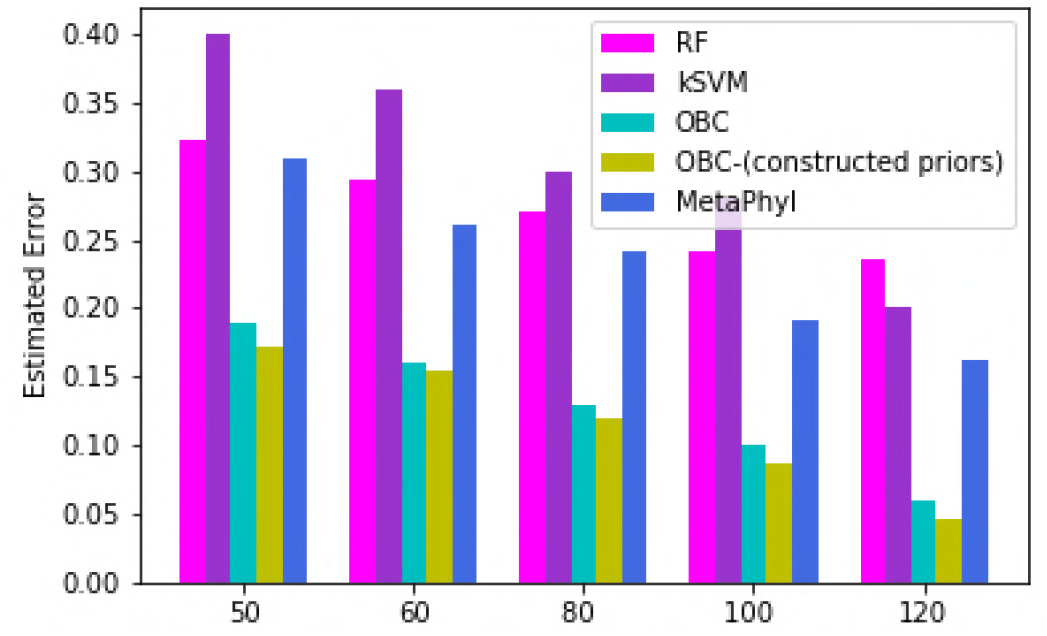
Average error rates for the various classification algorithms on the Twin Gut data set for varying training sample sizes.

## 3 Conclusion

In this paper, we presented a model-based Bayesian framework for the classification of metagenomic microbial abundance data. This approach was contrasted to state-of-the art metagenomic classification algorithms, including Random Forests and the phylogeny-based Metaphyl algorithm. The advantages of the proposed OBC classification algorithm include being applicable in the absence of phylogenetic information and in the presence of multiple classes and small ratios of sample size to dimensionality. The proposed classification method showed promising results in a comprehensive set of numerical experiments, particularly when the ratio of sample size to dimensionality is small.

## Acknowledgements

This work was supported by Texas A&M Engineering’s Strategic Initiative Seed program.

